# A Kaspar Hauser experiment for innateness of numerical cognition

**DOI:** 10.1101/2023.12.06.570352

**Authors:** Elena Lorenzi, Matilde Perrino, Andrea Messina, Mirko Zanon, Giorgio Vallortigara

## Abstract

Whether non-symbolic encoding of quantity is predisposed at birth with dedicated hard-wired neural circuits is debated. Here we presented newly-hatched visually naive chicks with stimuli (flashing dots) of either identical or different numerousness (with a ratio 1:3) with their continuous physical appearance (size, contour length, density, convex hull) randomly changing. Chicks spontaneously tell apart the stimuli on the basis of the number of elements. Upon presentation of either fixed or changing numerousness chicks also showed different expression of early gene *c-fos* in the visual Wulst, the hippocampal formation, the intermediate medial mesopallium, and the caudal part of the nidopallium caudolaterale. The results support the hypothesis that the ability to discriminate quantities does not require any specific instructive experience. Evidence for innateness of non-symbolic numerical cognition have implications for both neurobiology and philosophy of mathematics.

## Introduction

Quantity discrimination is well-attested in vertebrates^1–4^ and invertebrates^5^. Neurons selectively tuned to numerousness have been described in humans^6^, monkeys^6,7^ and crows^8^, and some evidence for number selectivity has been reported also in zebrafish^9–11^.

Artificial neural modeling suggests that units tuned to numerousness may emerge spontaneously in deep neural networks without any supervised learning^12^. However, the hypothesis that selectivity to numerousness is hardwired in the brain has been challenged^13^ and appears difficult to prove.

Early registration of number neurons showing adult-like properties has been reported in 8-12 day-old chicks^14^. However, although lacking any specific training on numerousness discrimination, these chicks have had plenty of opportunity to experience visual objects with their properties (including numerousness) during the days before testing. Similarly, some degree of experience was provided in all the behavioural studies performed so far in this precocial species (e.g., ^15–17^). Evidence for discrimination of the number of vertical bars in zebrafish larvae suffers a similar drawback: before tests, animals were reared extensively (for 4 days) in an environment with vertical bars in a simulation of natural landscape^18^. Besides, some of these studies lack proper control on the possibility that animals deal with discrimination using continuous physical variables that co-vary with numerousness (e.g. size, contour length, convex hull, etc.) rather than the number of elements *per se*. For instance, whereas chicks’ studies were well controlled for physical variables, in the experiments with zebrafish larvae the vertical bar stimuli were not equalized for e.g. spatial frequency (see for a well-controlled study in adult fish (archerfish)^19^). Thus, we do not know whether neuronal selectivity for number discrimination is present at the onset of life or it would require some specific experience in order to develop. Here we performed for the first time a «Kaspar Hauser experiment» for innateness of numerousness cognition by testing whether visually naïve newly-hatched chicks could detect spontaneusly a change in the number of elements in visual stimuli. We also investigated which brain regions would be involved in this discrimination.

## Results

According to the tradition of classical ethology^20^, the term “isolation” or “Kaspar Hauser experiment” refers to a proper control-rearing experiment, which would require testing visually inexperienced individuals for short durations to avoid discrimination being affected by learning through protracted exposure (note that the label came as only a generic reference to the historical character of Kaspar Hauser^21^). To meet with such requirements, we presented newly-hatched (dark incubated) visually naive chicks with stimuli (dots) with either identical or different (ratio 1:3) numerousness while continuous physical variables (size, contour length, density, convex hull, spatial frequency) randomly changed (Fig. 1 and see^19^ for details of the script for stimuli generation).

**Figure 1.**
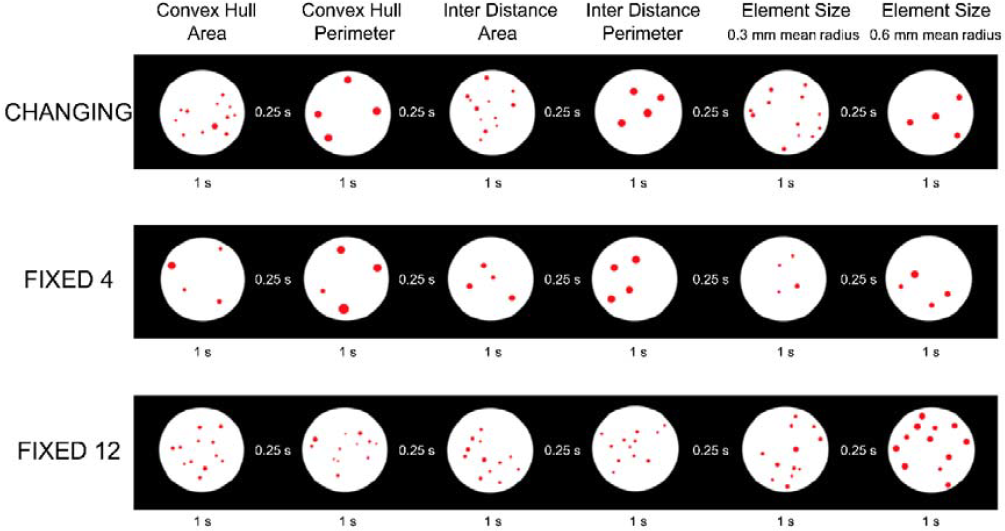
Experimental stimuli. Three different categories of stimuli sequences were used: Changing, Fixed 4 and Fixed 12. Columns display examples of stimuli (4 and 12) balanced from trial to trial for the different physical variables covarying with numerousness: Convex Hull and Area, Convex Hull and Perimeter, Inter Distance and Area, Inter Distance and Perimeter, single element Size 0.3 mm and 0.6 mm (see^22^ for details; the stimuli used were largely within the visual acuity capacity of newly-hatched chicks, which is in general much better than that of mammals^23^). Stimuli appeared for one second followed by a black screen with interstimulus intervals of 0.25 seconds, for a total sequence duration of 5 minutes.

We tested naïve chicks in a free choice test (Fig. 2a) for their spontaneous preference for the stimulus that showed a change in numerousness (from 4 to 12 dots and vice versa; *changing*) or for the stimulus that showed the same numerousness (*fixed*-4 dots, *fixed*-12 dots). We tested both males and females because a large body of evidence showed that males are more attracted than females by novelty in spontaneous preference tests and imprinting studies in chicks, including quantity estimation^24–28^. Thus, while we were testing whether both sexes were capable of detecting a change in number of elements, we also predicted that they could exhibit such an ability with different directions of choice (with a preference for change in numerousness in males and with a preference for the absence of change in numerousness in females).

**Figure 2.**
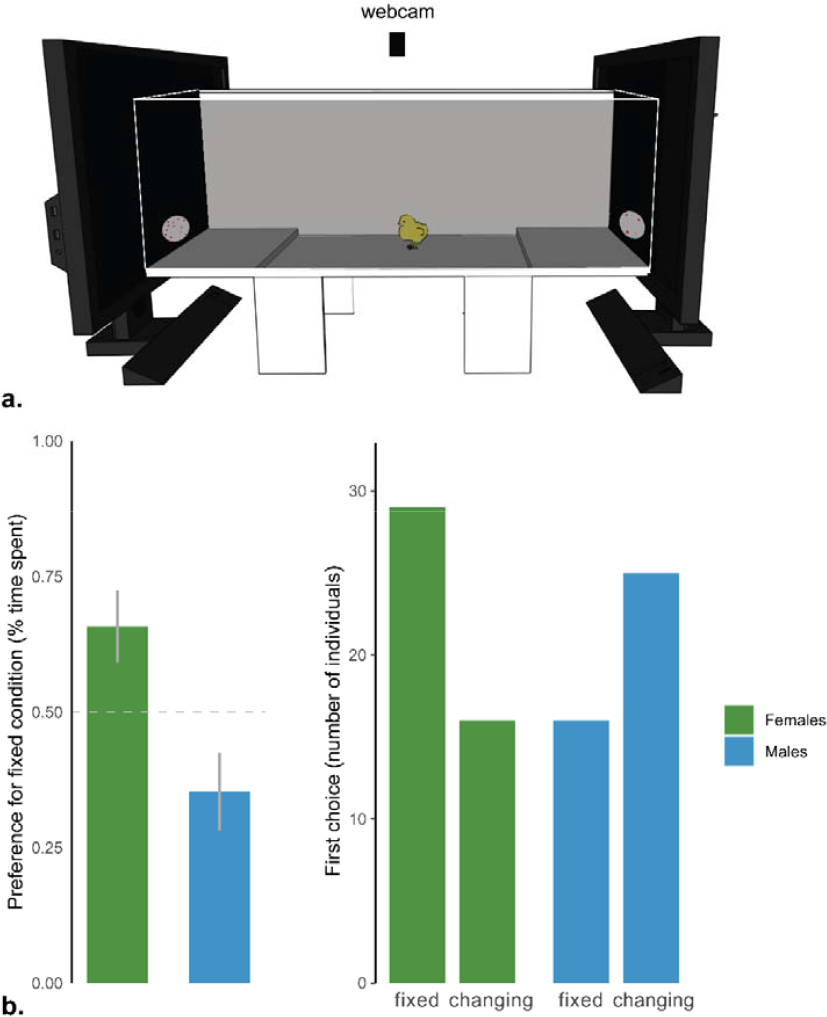
(a) Set up for behavioural testing. The two choice areas are delimited from a small step that chicks had to climb to access the zones near the two screens displaying the visual stimuli. For demonstrative purposes, one of the two long walls is depicted as translucent. (b) Percentages of time spent near the fixed numerousness stimulus condition (group means ± standard error are shown). Females (green) and males (blue) are shown separately. (c) First stimulus approached.

We analyzed the time spent close to the fixed condition as time ratio, considering the total time spent close to each stimulus across the 5 minutes:

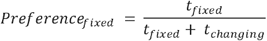

A permutation ANOVA with two factors (fix condition and sex; *aovp* R package was performed. The analysis showed a clear effect of sex (permutation p<<0.01) but no contribution of fixed condition or factors interaction.

It appeared that chicks spontaneously discriminated the two stimuli: females spent most time close to the fixed stimulus (t_(44)_=2.37, p=0.02, 95% CI: 0.52 – 0.79, Cohen’s d=0.35, N=45), males close to the changing stimulus (t_(40)_=-2.05, p=0.05, 95% CI: 0.21 – 0.50, Cohen’s d=0.32, N=41) (Fig. 2b).

First choices (Fig. 2c) showed the same pattern of results. A generalized model with binomial family (*glm* R package) showed only an effect of sex (i.e., the model with lower AIC=117.42 was the one including sex only).

Lumping together the two fixed conditions and performing a X^2^ test an effect of sex was apparent X^2^(1) =5.56 p=0.02, with female choosing more often the fixed numerousness and males the changing numerousness.

We then performed a second experiment in which separate groups of chicks were exposed to either the changing or the fixed stimulus of either number (Fig. 3a). We then measured *c-fos* expression in several brain regions to study which areas were mostly activated during the presentation of stimuli. We used only one sex for this experiment because we were interested in the ability to tell apart the two stimuli irrespective of the direction of choice, thus reducing the number of sacrificed animals. We selected females because they showed the clearest discrimination in behaviour (see Supplementary).

**Figure 3.**
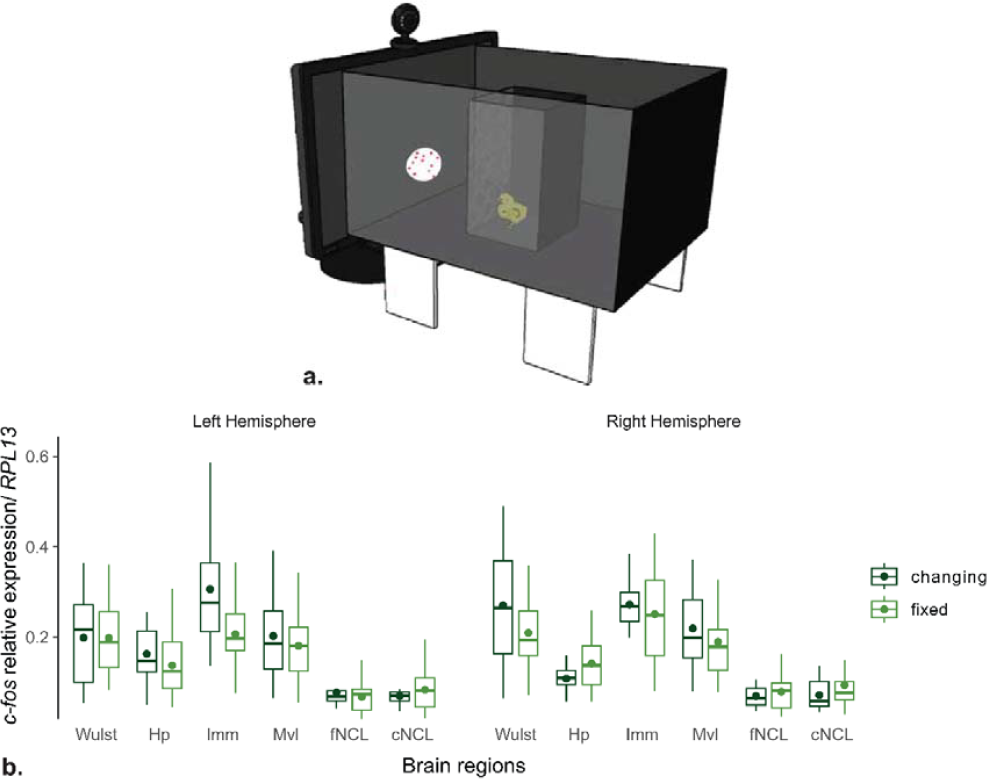
(a) Experimental apparatus for the neurobiology experiment. Schematic representation of the apparatus used during visual stimulation. The chick is represented within a small black box with a black metal grid on the side facing the video screen where the stimulus is presented. For demonstrative purposes, one of the two long walls is depicted as translucent. (b) Relative *c-fos* expression in different brain regions is shown separately for the fixed (light green) and changing (dark green) numerousness conditions in the left and right hemispheres. Boxplots represent the mean (dot) and the median (line), interquartile range (IQR, box) and maximum and minimum values within 1.5 ^*^ IQR (whiskers). Hp – hippocampus, IMM – intermediate medial mesopallium, MVL – ventrolateral mesopallium, fNCL – frontal nidopallium caudolateral, cNCL – caudal nidopallium caudolateral.

The brain areas of interest were selected on the basis of previous literature. In particular, we looked at the hippocampus because this is involved in response to novelty^29,30^ and we looked at plausible areas that are connected with the hippocampus but could be involved in processes more specific for the kind of information available here, namely change in numerousness. Thus, we selected the Nidopallium caudolaterale (NCL) because in this area number neurons have been reported in both chicks^14^ and adult crows^8^, and the Mesopallium, in particular the Mesopallium ventrolaterale (Mvl) and the Intermediate Medial Mesoplallium (IMM), which are associative regions in the avian brain that could be considered similar to temporal/parietal regions given of their connections with primary sensory areas and NCL (considered analogous of prefrontal cortex^33,34^). Also, the IMM has been described as partly involved in the initial formation of the recognition memory for imprinting.^31,32^ Moreover, given that there is mounting evidence that early visual areas could play a role in number perception (see reviews^2,35^) we also looked at the visual Wulst, which contains the avian equivalent of the mammalian primary visual cortex^33^.

Results are shown in Fig. 3b. A general permutation ANOVA with experimental groups (changing, fixed-4, fixed-12) as a between-subjects factor and areas (W, Hp, IMM, MVL and f/cNCL) and hemisphere as within-subjects factor was performed. We found clear effects of areas (permutation p<<0.01), hemispheres (permutation p<<0.01) and their interaction (permutation p=0.02), and most interestingly an interaction between area and group (permutation p=0.03). No significant heterogeneity was associated with the two numerousness in the fixed condition (fixed-4 and fixed-12): a permutation ANOVA with area, hemisphere and control group -2 levels: fix4 and fix12-as factor does not report any significant difference between control groups (permutation p=0.56; also, all interactions with condition factor are not significant). Thus, we merged these conditions and compared directly changing *vs*. fixed.

We found a significant effect of the interaction between experimental group and area (permutation p=0.02). We analyzed each area separately in order to investigate the specific contribution of each brain region.

In the Wulst there was a significant effect of hemisphere (permutation p=0.01) and an interaction of experimental group (fixed *vs*. changing) with hemisphere (permutation p=0.02): higher *c-fos* expression for changing than for fixed stimulus was observed in the right Wulst.

In the IMM a slight tendency for a hemisphere contribution (permutation p=0.08) and for the experimental condition (permutation p=0.13) was apparent. The left IMM was tendentially more active (changing vs. fix dt = 2.5, df = 15.7, p-value = 0.02, assuming equal variances).

In the hippocampal formation there was an effect of hemisphere (permutation p<0.01) and an interaction hemisphere with group (permutation p=0.04), with an activation following opposite directions in the two hemispheres as a function of the two stimulus conditions.

Finally, in the caudal part of the NCL we found an effect of group condition (permutation p=0.05) with higher activation in the changing than in the fixed condition irrespective of the hemisphere.

No significant effects were apparent in MVL and fNCL.

Interestingly, in the Wulst and possibly in the IMM *c-fos* expression was up-regulated in the changing numerousness condition, whereas it was down-regulated in the caudal NCL. This might be due to different activity of excitatory or inhibitory neurons in different areas that, however, given the methods used for the present experiment (qPCR) could not be further disentangled.

## Discussion

Chicks proved capable to spontaneusly discriminate the stimuli with changing and fixed numerousness. Their preference was immediately apparent at first choice, as well as in the overall percentage of time spent close to one or other of the two stimuli. As predicted, the two sexes expressed their preferences in different directions – for fixed numerousness in females and for changing numerousness in males – as noted (Introduction) it is well-documented in this species that males are more attracted by novelty^24–28^.

We identified some specific parts of the brain that showed a selective response to numerousness by measuring in separate groups of chicks’ early gene expression of *c-fos* following brief exposure to one or other type of stimulus. We found changes in *c-fos* expression in the hippocampal formation. Although hippocampal involvement in arithmetic in humans has been reported at the level of single cell activity^36^, recording in crows has not shown the presence of number neurons in the hippocampal formation^37^. Thus, it seems likely that hippocampal activation is mainly associated with response to novelty (which is greater for the changing than for the fixed stimulus because the former conveys changes in numerousness *per se* other than changes in continuous physical quantities). The kind of extra-novelty which is at play here – i.e. in the number of elements– could be detected mainly in the visual Wulst and in the NCL. Evidence that early visual areas could be implicated in number cognition has been suggested recently by work in both humans^38^ and zebrafish (^10,11^ and see for a review^35^); the NCL, on the other hand, is the region in which number neurons have been found in both crows^8^ and chicks^14^. The possible involvement of the IMM, a highly associative area in the avian brain, in number cognition is novel and intriguing: if confirmed might suggest that, as in the primate brain, there could be multiple sites for number neurons in the avian brain, similar to parietal and prefrontal cortices areas which role in number cognition has been described for both monkeys and humans^1^.

The results provide evidence for predisposed mechanisms to deal with number in the vertebrate brain. It is important to stress the novelty of our results: previous work with young animals, either altricial (e.g., human infants) or precocial (e.g., chicks) was always conducted with organisms tested after several days of life and extended visual experience (reviewed in^1,2^). Our chicks did not have any opportunity to estimate the number of visual objects before testing, and at test their ability showed up immediately at first choice. Thus, the role played by any kind of specific instructional experience here can be discarded, with the noteworthy implication for both cognitive neurobiology and the philosophy of mathematics that sensitivity to numerousness might be innately predisposed in the vertebrate brain.

## Acknowledgments

This work was supported by funding from the European Research Council under the European Union’s Horizon 2020 research and innovation program (Grant Agreement 833504 SPANUMBRA).

## Author contributions

E.L. and G.V. designed research; E.L. and M.P. performed research; M.Z. M.P. E.L. G.V. analyzed data; M.P., M.Z. and E.L. developed the stimuli, E.L and G.V. wrote the manuscript, all authors reviewed the manuscript.

## Declaration of interest

The authors declare no conflict of interest.

---

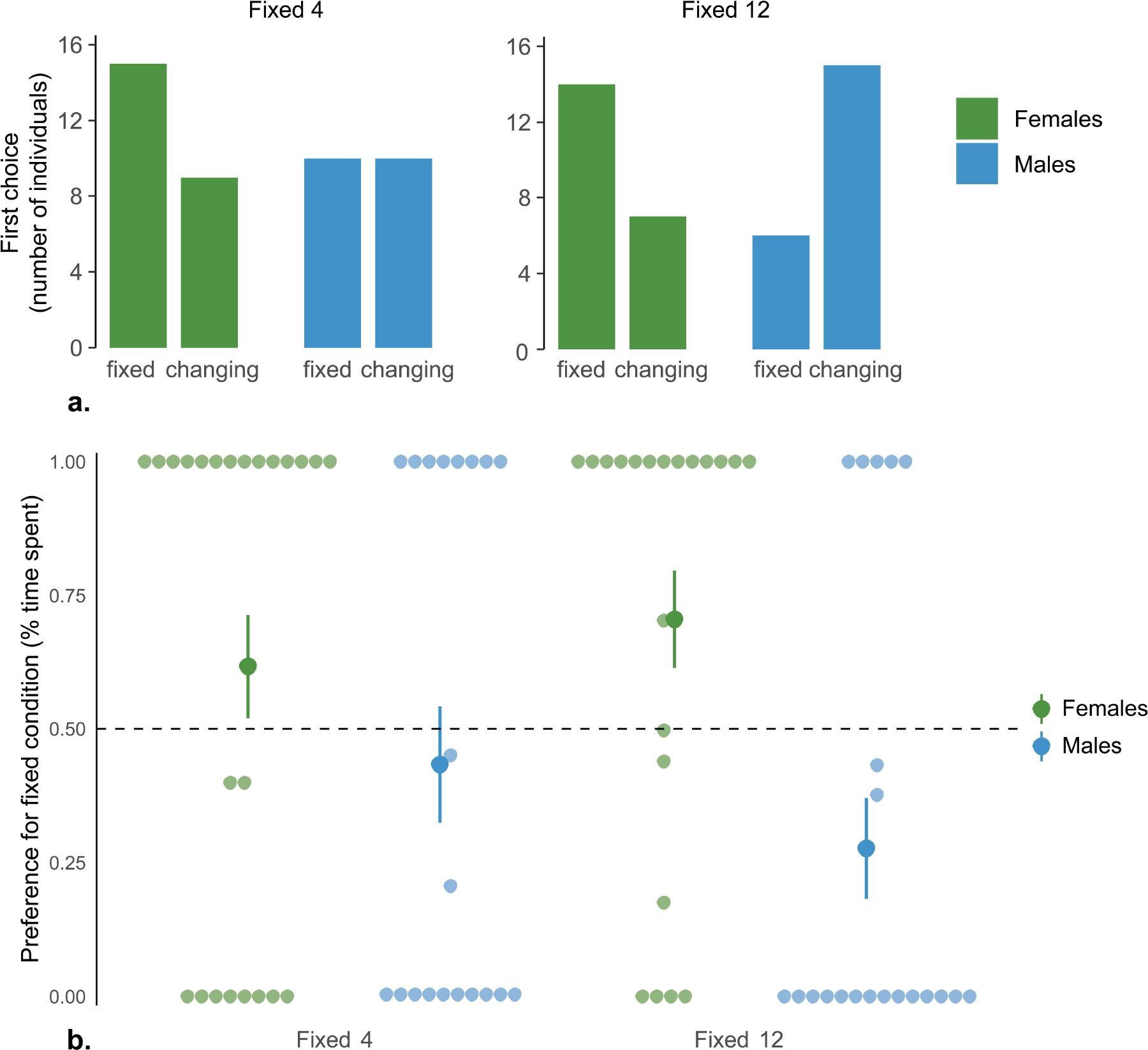

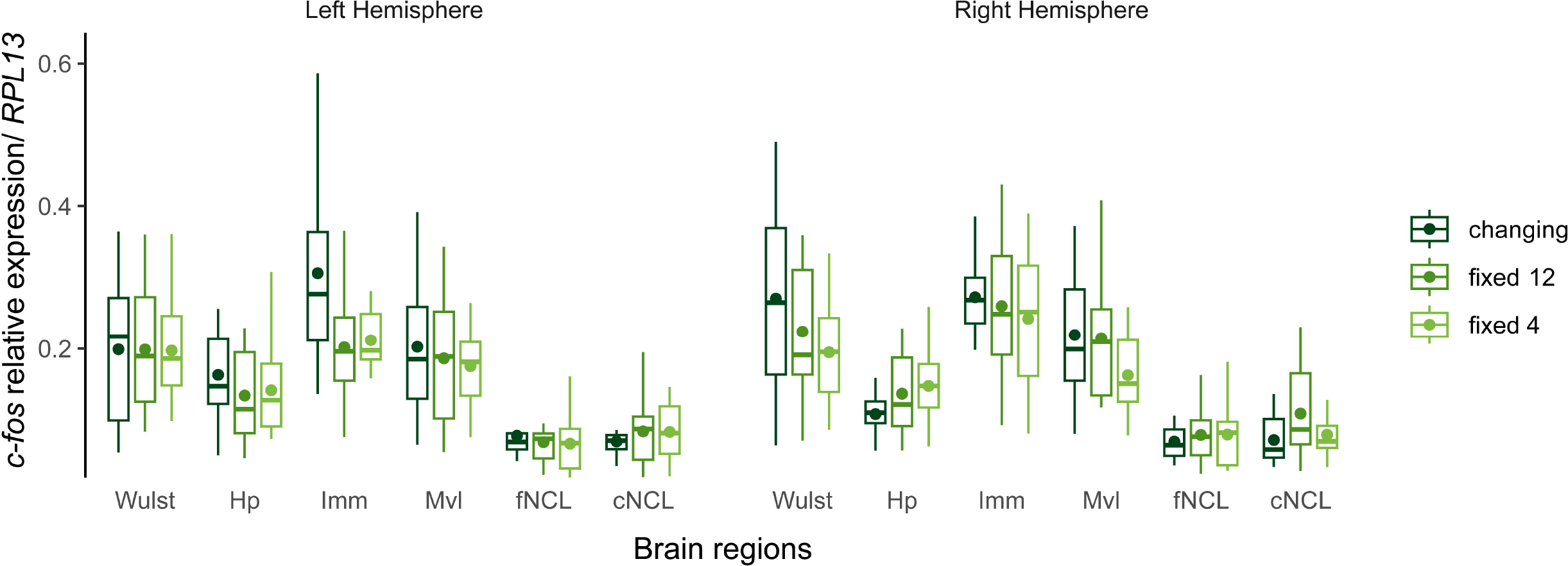

## Notes

https://doi.org/10.6084/m9.figshare.23748903.v1

### Competing Interest Statement

The authors have declared no competing interest.

