## Supplementary Information for "A Kaspar Hauser experiment for innateness of numerical cognition"

**Supplementary Materials**

Materials and Methods

### Animals

Newly-hatched domestic chicks (*Gallus gallus domesticus*) of the Aviagen Ross 308 strain were used. Eggs were obtained from a local commercial hatchery (Passirano, Brescia (BS) – Italy) and incubated in the dark in our laboratory at standard temperature and humidity until two days before hatching. Eggs were then moved to a Marans P140TU-P210TU incubator near the experimental setup, where they were incubated in total darkness at a constant temperature of 37.7°, with 60% humidity until hatching. The same conditions were maintained after hatching until testing. In order to avoid any visual experience before the test, all procedures were performed in complete darkness.

### Ethical statement

The experiments comply with all the applicable European Union and Italian laws and guidelines for animals’ care and use. All the experimental procedures were approved by the Ethical Committee of the University of Trento OPBA and by the Italian Health Ministry (permit number 791/2019).

### Visual stimuli

The rationale behind the visual stimulation used in numerical change detection procedure is inspired by the work of Starr et al, 2013 with human infants^1^. In our study, we used two numerousness, 4 and 12, represented by red dots inside a circular white arena of radius 1.5 cm (see Figure 1 main text). Stimuli were presented consecutively using the Psychophysics Toolbox Version 3 package in MATLAB R2019b for 1 second, followed by a black screen as an interstimulus interval of 0.25 seconds, for a total presentation time of 5 minutes. We used three different types of stimulus sequences (see Figure 1 main text): a sequence that showed the two numerousness in alternation (Changing) and two sequences with only one number of dots, either four or twelve (Fixed 4 and Fixed 12).

A ratio 1:3 was used as in previous studies in the same species (e.g.^2–4^) and in electrophysiological experiments in other bird species ^5^.

To counterbalance and control for the continuous stimulus variables co-varying with numerousness, stimuli were created in pairs (4 and 12) according to a dedicated script ^6^. Within each pair different combinations of continuous variables were controlled: Convex Hull and Area, Convex Hull and Perimeter, Inter Distance and Area, Inter Distance and Perimeter, single element Size of mean radius 0.3 mm and 0.6 mm. Ten pairs of stimuli with different spatial arrangements were generated within each of these categories. The values of convex hull, inter distance, area and perimeter were maintained the same across all their possible combinations; for example, stimuli with convex hull and perimeter controlled and inter distance and perimeter controlled had the same value for perimeter. Specifically, the controls were executed for each pair within each of the categories of controls ^6^. The sequences of the Change condition consisted of 20 pairs of counterbalanced stimuli for each of the six control categories randomly intermingled (see Figure 1), while the sequences of the Fixed conditions consisted only of one stimulus for each of the 20 couples (4 or 12) shown twice. Spatial frequencies were also counterbalanced in the script ^7^.

### Behavioural experiment

This experiment aimed at assessing chicks’ ability to spontaneously discriminate between the two sequences of stimuli, one with a change in the number of dots (Changing) and another with a fixed number of dots (Fixed) in a very short time and without any previous visual experience. Each chick observes the two streams, one of which alternates between the two numerousness, while the other maintains the same numerousness fixed though changing dot size and arrangement in space (above).

### Behavioural subjects

Eighty-six animals were used (45 females). Immediately after the test procedure, animals were caged in group in the animal house facility with water and food ad libitum, and shortly afterward donated to local farmers.

### Behavioural experimental apparatus

The apparatus (see Figure 2a main text) consisted of a white corridor (85 x 30 x 30 cm) with two opposite high-frequency video screens (ASUS MG248Q, 24’’, 120 Hz). It was subdivided into three areas: a central one (45 cm long) and two lateral ones (choice areas: 20 cm long each), which were divided by a small step (1.5 cm high) from the central area. A video-camera was placed above the apparatus and recorded the animals’ behaviour. Each screen presented a different sequence of stimuli: one showing always the same numerousness (Fixed) and the other showing the change in numerousness (Changing). The luminosity of each screen was calibrated and maintained equal.

### Behavioural testing

After hatching, each animal was taken singly from the incubator in darkness and transported to the experimental room into a dark box. It was placed in the middle of the central area facing one of the two long walls of the apparatus. Then, the experimenter started the stimulus sequence presentation. All chicks saw the Changing sequence on one screen contrasted by one of the two Fixed sequences on the opposite screen. Fixed sequences (4 and 12 number of dots) were counterbalanced between-subjects (fixed 4: females = 24, males = 20; fixed 12: females = 21, males = 21). We counterbalanced between subjects both the side of stimulus presentation (left, right), as well as the orientation of the chick’s beak at the beginning of the test, in order to control for any side bias. Data from animals that did not move from the central area for the whole duration of the experiment were discarded from further analyses.

### Behavioural statistical analyses

For the behavioural analysis we evaluated both the time ratio spent close to the fixed condition and the very first choice.

For the time ratio, we considered the total time spent close to each stimulus across the 5 minutes presentation, and calculated the percentage of this time close to the fixed condition:

$$Preference_{fixed} = \frac{t_{fixed}}{t_{fixed}+ t_{changing}}$$

We analyzed this variable with an exact permutation ANOVA (*aovp* function in R) with sex (2 levels: females, males) and experimental condition (2 levels: fixed 4 or fixed 12) as factors. Given the nature of the test used p-values oscillates at each permutation repetition; here we reported the most frequent p-value obtained from multiple executions of each test.

For the first-choice analysis we run both a generalized linear model with binomial family (*glm* function in R) - fitting models of different complexity and evaluating the best fit following Akaike information criterion (model with minimum AIC) -, and a chi squared test with only the sex factor (as indicated by previous generalized model). The post-hoc analysis at the single sex level was performed with a binomial test.

### Neurobiological experiment

The goal of this experiment was to investigate the main areas of the chick’s telencephalon involved in the perception of numerousness change. Each chick was exposed to one of the three sequences (Changing, Fixed 4 and Fixed 12) using the same stimuli as in the previous experiment. Neural activity was assessed by RT-qPCR quantification of different Immediate Early Genes relative expression as marker of synaptic plasticity in the main telencephalic areas of the chick’s brain. IEGs estimated: *c-fos*. Expression was normalized across subjects through the quantification of the reference gene *RPL13* involved in basic cell functioning as an internal control.

### Neurobiological Subjects

A total of 45 chicks were used (all females) for this neurobiological experiment. A group of chicks was exposed to the Changing sequence (N=15), a group was exposed to the Fixed 4 sequence (N=15) and a third group was exposed to the Fixed 12 sequence (N=15).

### Neurobiological experimental apparatus

The apparatus consisted of a black wood box (37.3 cm x 49.5 cm x 28.3 cm) with an open side where a video screen 30 cm x 53 cm (ASUS MG248Q, 24’’, 120 Hz) was inserted (Figure 3 main text). A small black plastic box (14.5 cm x 14.5 cm x 23.3 cm) was placed inside the main container at a distance of 35 cm and it was used to constrain the chick during the visual exposure. The side of this box in front of the video screen was made of a black metal grid, which thus enabled the view of the stimuli. The distance of the box was calculated in relation to a stimulus subtended visual angle of ~ 4° (Weller 2009, Meyer & May 1973), obtained through the formula:

$$\alpha= 2 arctan(|ba| \div d)$$

A webcam was placed centrally above the video screen to enable frontal recording of chick’s behaviour, all the video recordings were kept and saved for subsequent off-line analyses.

### Neurobiological testing

After hatching, each animal was taken singly from the incubator in darkness and transported to the experimental room into a dark box. The animal was placed in the exposure box with the beak facing the video screen. After the positioning, one of the three stimulus sequences started and the animal was free to move inside the box for the whole stimulation. In order to sample only subject that had paid attention to the stimulus, we excluded from further analyses animals that fell asleep during the experiment or that looked at the stimulus for less than 3.5 minutes. After the visual stimulation, each subject was carefully placed back into the same incubator, in a single compartment (cardboard box 11 cm x 11 cm x 25 cm). This procedure was performed in order to keep the auditory environment as it was before the exposure.

### Quantitative reverse transcription PCR (RT- qPCR^8^)

### Tissue acquisition

After 30 minutes from the visual stimulation, each chick received an overdose with an intramuscular injection of 0.05 ml Ketamine/Xylazine Solution (1:1 Ketamine 10 mg/ml + Xylazine 2 mg/ml) per 10 gr of body weight. After 5 minutes, the animal became unresponsive and the head was severed and immediately placed in ice. The skull was fixed on a stereotaxic head holder and it was surrounded by ice to prevent the deterioration of RNA. The skin and skull were partially opened and the dorsal surface of the brain exposed. Six coronal cuts of 1mm thickness were made from left to right at precise coordinates in order to obtain different slices on the rostro-caudal axis from plate A(nterior) 12.8 to plate A4.8 ^9^) With surgical spatula and scoop, the slices were removed from brain one after the other following a rostro-caudal order. The main areas of interest were: the Wulst, the Intermediate Medial Mesopallium (IMM), the Ventrolateral Mesopallium (MVL), the Hippocampus (Hp), the frontal Nidopallium Caudolateral (fNCL and the caudal Nidopallium Caudolateral cNCL). The areas were selected respectively from slices corresponding to Kuenzel and Masson’s^9^ atlas plates A12.6 for Wulst, A8.8 for IMM and MVL, A7.0 for Hp, A5.4 for fNCL and A4.8 for cNCL. Areas were cut off from brain slices and collected in a separate 1.5 ml eppendorf, which was immediately stored at -20°C.

### RNA extraction

From each sample, the total RNA was extracted with The RNeasy Mini Kit (QIAGEN Group) according to the manufacturer’s instructions. At this step, the Total RNA present in the cells was extracted. The tissue was disrupted and homogenized within a lysis buffer, then centrifuged and a volume of 70% ethanol was added to the lysate. The sample was moved to a spin column so that the RNA would bind to the filter membrane and then centrifuged. To eliminate any DNA molecule from the sample, a DNase I incubation mix was directly added to the RNeasy column membrane and incubated at 20°C for 15 minutes. A succession of washes with salts and ethanol and centrifuges was performed to purify the RNA solution. At the end, the RNA was eluted with 30 μL RNase-free water. A measure of RNA concentration and purity (A260/A280 and A260/A230 values) was obtained with the NanodropTM spectrophotometer (NanodropTM Onec; ThermoFisher Scientific, USA). Each sample was then stored at -20°C.

### Reverse transcriptase-polymerase chain reaction

In order to be able to quantify the expression levels of specific RNA, quantitative Reverse Transcriptase Polymerase Chain Reaction (qRT-PCR) was performed using a reverse transcriptase enzyme to produce single-stranded codifying RNA to single-stranded codifying DNA (cDNA) while preserving relative concentration ratios of different RNAs. For this procedure, SuperScriptTM VILOTM cDNA Synthesis Kit (Invitrogen, ThermoFisher Scientific, USA) was used following manufacturer’s instructions. For a final solution of 20 μL, 2 μL of enzymes, 4 μL of buffer solution and 14 μL of RNA were used. The 14 μL of RNA were normalized to a concentration 1μg/14μL when the RNA concentration. The cDNA was then diluted to a concentration of 2.2 ng/μL in order to get 10 ng of cDNA in 4.5 μL of solution. This reaction is performed with C100 TouchTM Thermal Cycler (Bio-Rad, USA).

### Quantitative Polymerase Chain Reaction

qPCR was used to quantify the expression of *c-fos* and *RPL13* by using specific forward and reverse primers (Table 1). In particular, qPCR works through the amplification of cDNA by DNA-polymerase chain reactions through thermal cycles and the consequent cDNA quantification at each cycle. The amplification is obtained through the exposure of reactant to repeated cycles of heating and cooling to permit DNA melting and enzyme-driven DNA replication, respectively at temperatures of 95°C and 60°C.

| Gene primer | Sequence |
| --- | --- |
| *c-fos* Fw | GTGTTCCTGGCAATATCGTG |
| *c-fos* Rw | TCAGACCACCTCAACAATGC |
| RPL13 Fw | TCGTGCTGGCAGAGGATTC |
| RPL13 Rw | TCGTCCGAGCAAACCTTTTG |

Table 1: Gene sequences of the primers (forward and reverse) used in the RTqPCR.

The quantification was recorded through a measure of fluorescence intensity emitted by the dye (SYBR Green) that intercalates in all double-stranded cDNA. The process of amplification of specific DNA products leads to an increase of fluorescence intensity measured at each cycle, being the fluorescence a function of the quantity of cDNA in a fixed amount of time. The compound is excited with blue light (λ max = 497 nm) and emits back green light (λ max = 520 nm). Moreover, at the end of each amplification the specific melting curve of each DNA molecule is determined as a quality control, since each cDNA in this case has a precise melting temperature, depending on the composition of amplicon bases. We used 5 μL of Master Mix, 0.5 μL of primers (forward + reverse) and 4.5 μL of cDNA, at a concentration of 2,2 ng/4.5 μL. The PowerUpTM SYBRTM Green Master Mix (Applied Biosystems, USA) included the SYBR Green dye. For each sample, three replicates were made to increase the precision of the measurement.

The reaction was performed with CFX96TM Real- Time System (Bio-Rad, USA). The output data consisted in quantification cycle (Cq) values, which represents the point in time across the 40 cycles (i.e. the number of cycles) when the fluorescence signal reached the threshold. The points outside the range of ±0.50 from the median value of each triplicate were excluded and the mean of the triplicates was made. For each sample, the Cq of *c-fos* was compared to the Cq of the reference gene RPL13 with the formula:

$$deltaCq=2^{(Cq cfos-Cq Rpl13)}$$

The expression of each gene of interest was normalized through the formula:

### Neurobiological statistical analyses

Outliers were evaluated with the IQR criterion (observations above q_0.75_ + 1.5⋅IQR or below q_0.25_ − 1.5⋅IQR; where q_0.25_ and q_0.75_ correspond to first and third quartile respectively, and IQR is the difference between the third and first quartile) for each area, hemisphere and group were removed. Outliers were considered random technical errors in either molecular biology procedures or in association with time viewing (chicks may move independently their eyes and use both frontal and lateral vision thus inter-individual differences in time looking at stimulus were expected).

For the statistical analysis we performed permutation ANOVA tests. Given the nature of the test used p-values oscillates at each permutation repetition; here we reported the most frequent p-value obtained from multiple executions of each test.

To simplify the overall analysis and focus on our experimental question we proceed like this:

1.	We run a first general permutation test on the relative cfos expression with hemisphere (2 levels: left, right), brain area (6 levels: Wulst, Imm, Hp, Mvl, cNCL, fNCL) as within factors and experimental condition (3 levels: changing, fixed 4, fixed 12) as between factor.

2.	Given an effect of experimental condition, we performed a focused analysis with fixed condition only (hemisphere, brain area as within factors and fixed condition - 2 levels: fixed 4, fixed 12- as between factor). Given the absence of difference between these two control conditions we merged them for the following analysis, to focus on our question of interest: the overall difference between changing and fixed conditions. Moreover, given a clear area differentiation (as expected), for the following we performed separated analysis for each brain area.

3.	Thus, separated permutation ANOVA tests were performed in each brain area with hemisphere (2 levels: left, right) as within factor and experimental condition (2 levels: changing and fixed) as between factor. Post-hoc analyses were performed with t-tests.

For each area, the numbers of experimental subjects (change and fix groups) analyzed are 15 (change) and 30 (fixed). Depending on outliers, the number of hemispheres for each area and group varies.

### Data availability

Data and code used for analysis are available at <https://doi.org/10.6084/m9.figshare.23748903.v1>

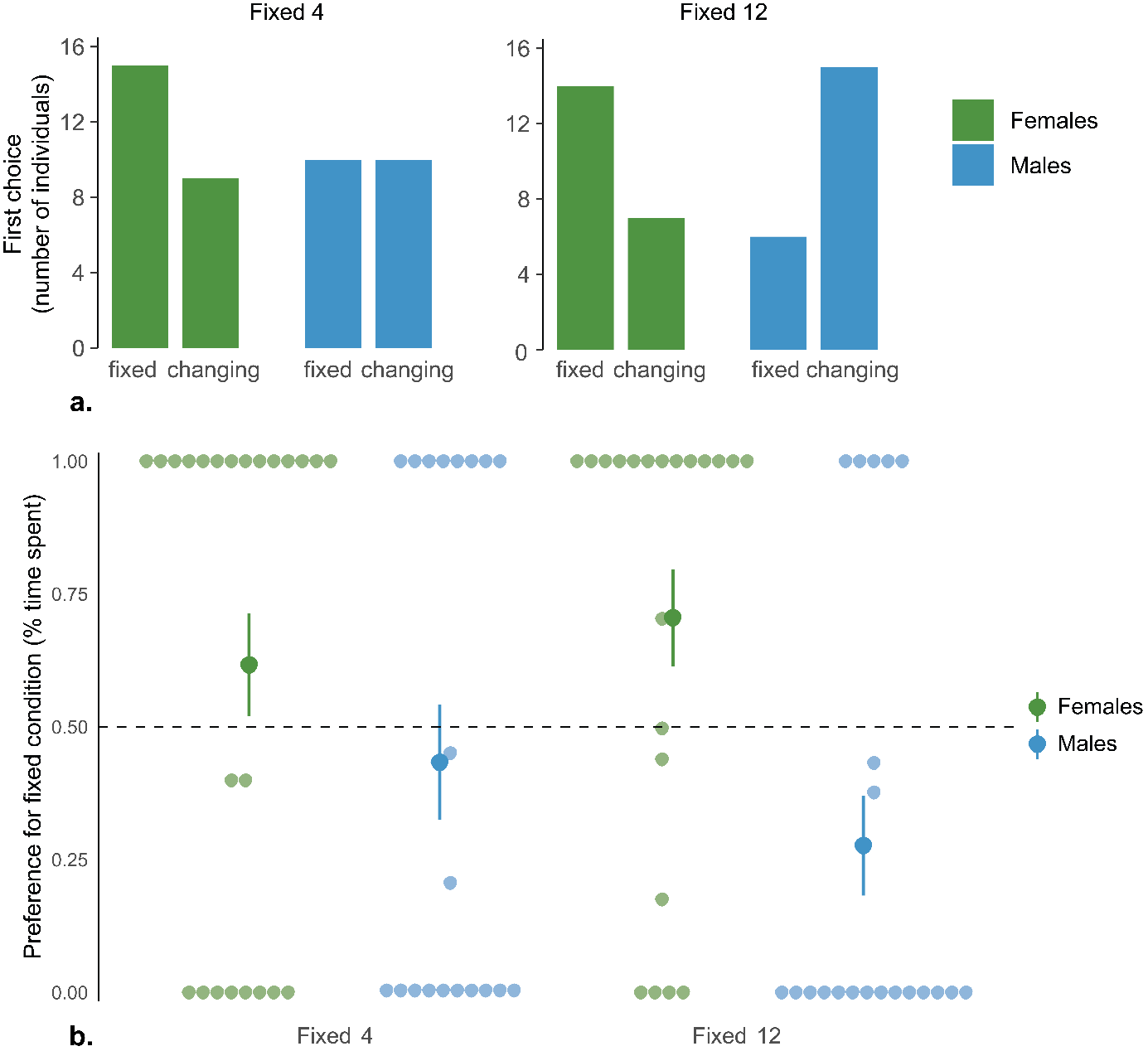

Figure 4 (a) First stimulus approached, for either fixed numerousness 4 or 12. Females (green) and males (blue) are shown separately. (b) Percentage of time spent near the fixed numerousness stimulus condition for either fixed numerousness 4 or 12 (individual values and group means ± standard error are shown).

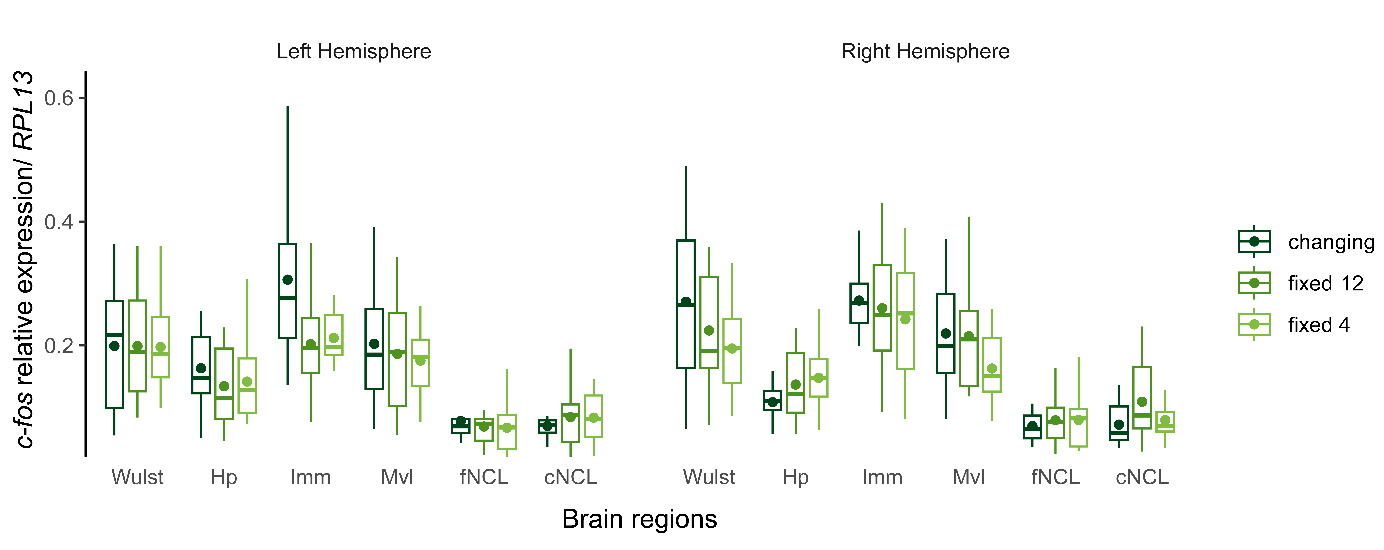

Figure 5 - *R*elative *c-fos* expression in different brain regions is shown separately for the changing (dark green), fixed 12 (light green) and fixed 4 (yellow) conditions in the left and right hemispheres. Boxplots represent the mean (dot) and the median (line), interquartile range (box) and maximum and minimum values whitin 1.5 * IQR (whiskers). Hp – hippocampus, IMM – intermediate medial mesopallium, MVL – ventrolateral mesopallium, fNCL – frontal nidopallium caudolateral, cNCL – caudal nidopallium caudolateral.
